# Assessing the Impact of Valve Sizing and Orientation on Bioprosthetic Pulmonary Valve Hemodynamics Using In Vitro 4D-Flow MRI

**DOI:** 10.1101/2024.05.09.593439

**Authors:** Nicole K. Schiavone, Priya J. Nair, Christopher J. Elkins, Doff B. McElhinney, Daniel B. Ennis, John K. Eaton, Alison L. Marsden

## Abstract

**Purpose:** Pulmonary valve replacement (PVR) using bioprosthetic valves is a common procedure performed in patients with repaired Tetralogy of Fallot and other conditions, but these valves frequently become dysfunctional within 15 years of implantation. Since PVR is often performed in adolescence, valves are typically oversized to account for somatic growth. However, the contribution of oversizing to early valve failure are not clearly understood. The purpose of this study was to explore the impact of valve sizing and orientation on local hemodynamics and valve performance.

**Methods:** Different valve sizes were represented by changing the cardiac output through a 25 mm bioprosthetic valve implanted in an idealized 3D-printed model of the right ventricular outflow tract (RVOT). The local hemodynamics at three valve sizes and two valve orientations were assessed using 4D-Flow MRI and high-speed camera imaging.

**Results:** Noticeable differences in jet asymmetry, the amount of recirculation, leaflet opening patterns, as well as the size and location of reversed flow regions were observed with varying valve sizes. Rotation of the valve resulted in drastic differences in reversed flow regions, but not forward flow.

**Conclusion:** Flow features observed in the oversized valve in this study have previously been correlated with calcification, hemolysis, and leaflet fatigue. Therefore, valve oversizing can negatively impact local hemodynamics and leaflet performance.

## 1 Introduction

Pulmonary valve replacement (PVR) is a common re-intervention procedure in patients with repaired Tetralogy of Fallot (ToF), typically indicated to treat pulmonary regurgitation (PR). For ToF patients, the first PVR is often performed in adolescence, with additional reinterventions often needed over a patient’s lifetime [1– 3]. After PVR, most patients experience significant reduction in right ventricle (RV) end-systolic and end-diastolic volumes, and the amount of PR is drastically reduced or eliminated entirely [4–6].

Bioprosthetic valves are among the most commonly used prostheses for PVR in ToF, since mechanical valves require long-term anti-coagulation therapy, which poses significant risks in children [7–10]. However, most bioprosthetic valves experience structural deterioration and dysfunction over time, often within 15 years of implantation [10, 11]. In ToF patients, these valves can fail early and unpredictably in as many as 30% of patients [12, 13]. Many studies have noted that younger age at the time of PVR is a predictor of shorter longevity of both bioprosthetic valves and valved conduits [7–9, 12, 14, 15]. However, within this younger cohort, it is currently not possible to predict which patients will experience earlier valve dysfunction. In addition, there are no standard surgical techniques proven to prolong bioprosthetic pulmonary valve durability.

Clinical observation reveals a wide range of RVOT anatomies within the ToF patient population [16–18]. Common anatomic variations include significant dilation of the main pulmonary artery (MPA) local to the valve, as well as variations in bifurcation angles, vessel lengths and diameters in the RVOT, pulmonary trunk and branch pulmonary arteries (PAs)[16]. Previous studies have shown that these anatomic features, as well as the placement of prosthetic valves, alter the hemodynamics of the RV and PAs from expected flow patterns seen in healthy anatomies [19–25]. Thus, understanding how bioprosthetic valve placement in ToF anatomies impacts the hemodynamics local to the valve may provide insights into valve function over time.

Valve sizing is a critical decision during surgery, especially in younger ToF patients undergoing PVR. It is possible that early valve failure may be due to somatic growth, which leads some surgeons to oversize the valve, placing a larger valve than necessitated by the patient’s native RVOT size. However, the effects of valve oversizing are not fully understood. Some studies of bioprosthetic pulmonary valves and RV to PA valved conduits have found that valve oversizing decreases longevity and is associated with shorter freedom from dysfunction [7, 8, 26]. In addition, a computational study examining three different sizes of valved conduits placed virtually in a healthy pediatric PA found that mean wall shear stress, an indicator for stenosis in these prostheses, was 40% higher at the anastomosis in simulations with an oversized conduit [27]. In studies using stentless aortic valves, valves that were matched with the true patient internal diameter had higher efficiency [28, 29]. However, recent studies have demonstrated that valve oversizing is not associated with significant changes in time to dysfunction or reintervention [14, 15, 30]. The effect of valve oversizing therefore remains an open clinical question.

Clinical 4D-Flow MRI, which uses phase-contrast imaging to obtain velocity fields, has been used in numerous studies examining flow in a range of cardiovascular diseases [31–37], including ToF, but the effects of valve placement or sizing have not been examined [6, 36, 38, 39]. To assess valve performance, it is often useful to capture the motion of the valve leaflets in addition to quantifying the surrounding velocity fields. High-speed imaging is used frequently to visualize the valve leaflets over the cardiac cycle and to calculate various performance metrics including effective orifice area and leaflet bending stress [40–42]. However, high-speed imaging studies have not often been done in conjunction with 4D-Flow velocity field measurements. In this study, we demonstrate the benefits of obtaining both velocity fields and high-speed imaging data for the same set of experiments.

## 2 Methods

All experiments in this study used the same 25mm Epic valve (St. Jude Medical, Abbott). By changing the cardiac output (CO) through the same size valve, we reproduce the clinical scenario of changing valve sizing. CO generally aligns with patient size: smaller patients have lower CO. Thus, when a surgeon places a large valve in a smaller patient, they are placing the valve in an environment with lower flow than if it had been placed in an appropriately sized patient. The flow loop was tuned to three different COs: 2 L/min, 3.5 L/min, and 5 L/min. The 3.5 L/min case represents the typical flow for a patient with an appropriately sized 25mm valve. The 2 L/min case corresponds to valve oversizing in a clinical setting and the 5 L/min case corresponds to a patient outgrowing their valve.

The dilated MPA model described in Schiavone *et al*. was used in this study[22]. The model and flow loop were used to obtain 4D-Flow MRI velocity data and to capture valve leaflet images using high-speed imaging in separate benchtop experiments.

The working fluid for the flow loop was a blood analog fluid of 60% water and 40% glycerin, with a density of 1.1 g/cm3 and a viscosity of 3.9 cP at room temperature. Gadolinium was added to the blood-analog fluid to increase the signal intensity during the 4D-Flow MRI scan. The inflow to the right ventricle (RV) box was measured via an ultrasonic flow probe while pressure transducers (Micro-Tip SPR-350, Millar) were used to measure pressures proximal and distal to the valve. Various components of the flow loop were adjusted from the 3.5 L/min baseline to obtain the different cardiac outputs. For the 2 L/min case, the duration of the sine squared waveform driving the pneumatic system was decreased. For the 5 L/min case, both the duration and the amplitude of the sine squared waveform were increased. These changes affected the forward air flow into the RV box, which in turn affected the pulsatile flow rate throughout the flow loop. However, while these changes allowed us to capture the correct net flow rate, they adversely affected the pressure waveforms. To account for these changes, we realigned the venting of the pneumatic box with the sine squared waveform to replicate similar dynamics in the driving system across all cases. In addition, we increased the downstream capacitance for the 5 L/min case and decreased the capacitance for the 2 L/min case. In order to maintain a similar pressure waveform shape across all cases, the length of systole varied slightly depending on the cardiac output. Systole was approximately 33% of the cardiac cycle for the 2 L/min case, 38% of the cardiac for the 3.5 L/min case, and 41% of the cycle for the 5 L/min case. The adjustments to the system resulted in physiological flow and pressure waveforms for each case, as seen in Figure 1. Overall, we generated similar dynamics for each cardiac output, allowing for the most direct possible comparison between all three cases. Physiological flow and pressure waveforms were obtained (Figure 1).

**Fig. 1.**
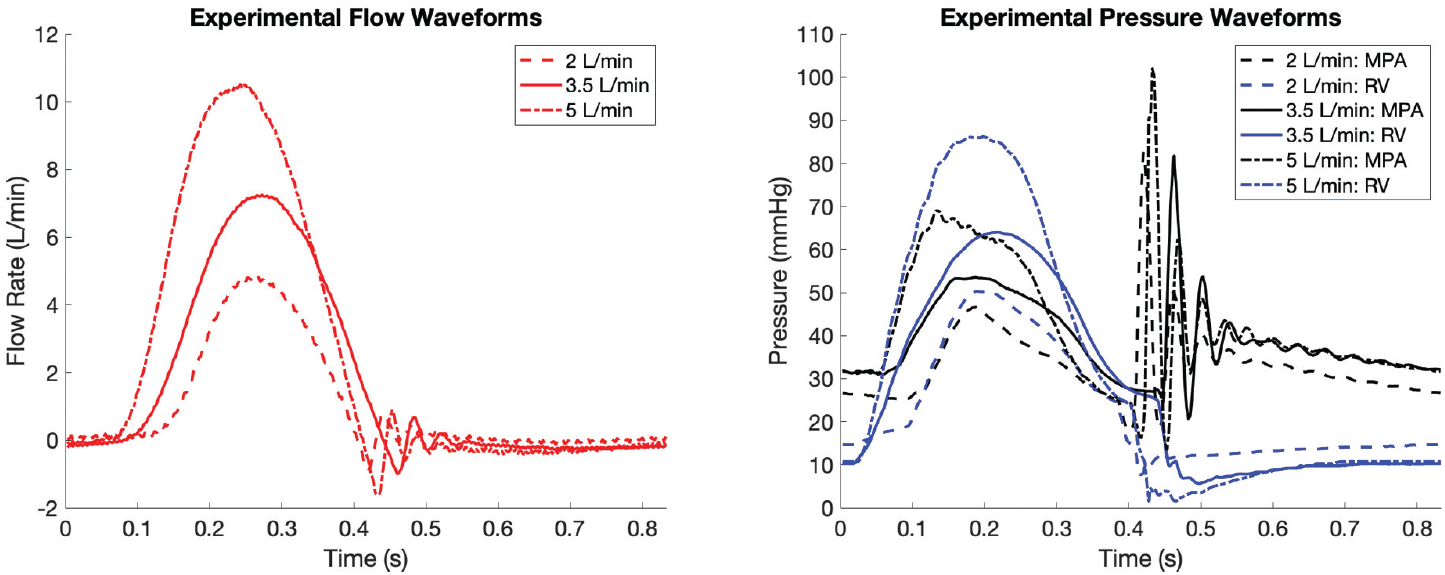
Experimental flow (left) and pressure (right) waveforms in the model for each of the three CO conditions. Flow waveforms were measured in one of the outlet PA branches. Flow and pressure data were taken simultaneously for each case.

### 2.1 4D-Flow MRI Experiments

To analyze the effects of CO and valve orientation, we conducted five experiments. We ran the flow loop with COs of 2 L/min, 3.5 L/min, and 5 L/min with the valve in the native orientation – abbreviated as *V*_N,2_, *V*_N,3.5_, and *V*_N,5_. For the rotated orientation, we only ran the 2 L/min and 3.5 L/min cases (abbreviated as *V*_R,2_ and *V*_R,3.5_), since 2 L/min represents the more clinically relevant case of valve oversizing and 3.5 L/min represents a correctly-sized valve.

The scan procedure and sequence settings for the MRI were nearly identical across all experiments and are outlined in Schiavone *et al*. [22]. The slice direction was sagittal, with a spatial resolution of 0.9 mm isotropic and scan time of 21 minutes. The temporal resolution was 77 milliseconds, which allowed for direct measurement of 10 phases; these were used to interpolate to 20 total phases in the cardiac cycle. Therefore, for each scan, we report phase-locked, 3D, three-component, time-averaged velocity fields for 20 phases of the cardiac cycle. The velocity encoding (VENC) was the only parameter that was changed, since changing the CO results in different maximum velocities for all three cases. VENC is a parameter that specifies the maximum velocity that can be encoded without any phase aliasing. The VENC in each case was set to the same value in the sagittal, axial, and coronal directions. The VENC was set at 150 cm/s for the 2 L/min case, 250 cm/s for the 3.5 L/min case, and 350 cm/s for the 5 L/min case.

We calculated the signal-to-noise ratio (SNR) and velocity uncertainty for each case. The expected uncertainty of 4D-Flow MRI velocity measurements is given by

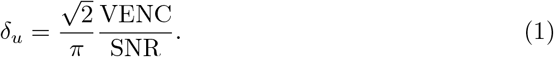

We report the velocity uncertainty as the percent of the mean velocity through the valve opening at peak systole. The average SNR for the 3.5 L/min cases was 21, resulting in an uncertainty of 6.3%. For the 5 L/min case, the SNR was 18, resulting in an uncertainty of 7.6%. The average SNR in the two 2 L/min cases was 19.5, resulting in an uncertainty of 6.2%.

To quantify the streamwise momentum and secondary flow strength of the flow fields, we calculated the integral metrics *I*_1_ and *I*_2_ for each case over the cardiac cycle defined by

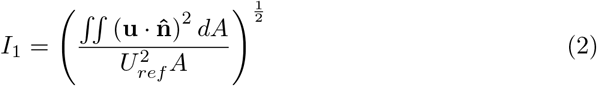

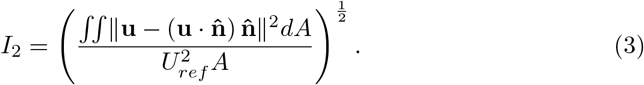

Here, **u** is the velocity vector, 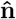 is the unit vector normal to the cross-section, and *A* is the local cross-sectional area of the geometry at the plane where *I*_1_ and *I*_2_ are evaluated. *I*_1_ and *I*_2_ were calculated for the axial cross-sections at *x/D* = 0.25 and *x/D* = 0.5. For each case, *U*_*ref*_ was the mean speed in the RVOT immediately upstream of the valve at peak systole for the reference area.

### 2.2 High-Speed Imaging Experiments

For the high-speed imaging experiments, the flow loop was assembled on the benchtop exactly as it was in the MRI scanner. Though the high-speed imaging and the 4D-Flow MRI experiments are not conducted simultaneously, they both use and record the same trigger signal for the pulsatile cardiac cycle. Thus, we can precisely align the data from the two methods. We obtained high-speed images of the valve leaflets for the three COs with the valve in the native and rotated orientation for each case.

In order to have clear optical access to the valve, we printed a new downstream component for the MPA and branch PAs of the model. The internal geometry of the model was identical, but we reduced the wall thickness to 2mm and polished the surface to remove distortions due to the printing process. To eliminate refraction from the surrounding air to the working fluid in the model, we filled an external box (black arrow in Figure 2) with the working fluid to submerge the model. We captured images of the valve leaflets, head-on from downstream, using a Phantom V2012 Ultrahigh-speed camera with a 105mm Nikkor lens.

**Fig. 2.**
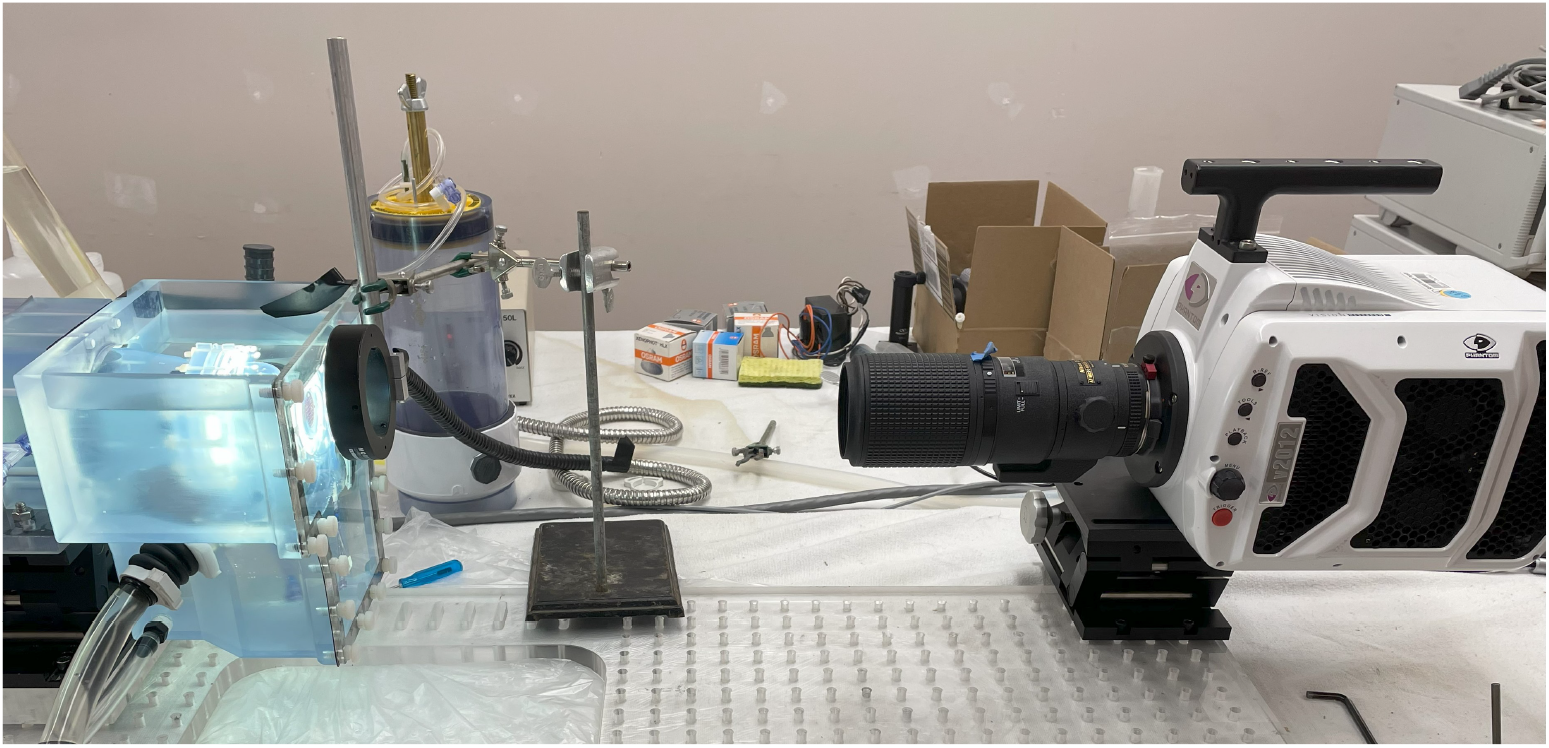
The experimental flow loop assembled on the benchtop for high-speed imaging. The Phantom V2012 Ultrahigh-speed camera on the right was aligned head-on with the valve in the model in order to capture images of the leaflet motion. The model was submerged in the working fluid to eliminate refraction effects.

We acquired images at a frame rate of 1500 Hertz with a 500 microsecond exposure to capture the instantaneous valve leaflet motion. For a heart rate of 72 bpm, this frame rate resulted in approximately 409 frames during systole for the 2 L/min case, 476 frames during systole for the 3.5 L/min case, and 514 frames during systole for the 5 L/min case. We used an f-stop of 11 to allow for a large enough depth of field to keep the leaflets in focus as they open. A ring light was placed on the outside of the submersion box, positioned with the valve centered in the ring, to provide lighting for high quality imaging.

We used a calibration target with a grid of 69 evenly spaced points that fit into the valve-holder component of the model. The target was set at the location where the leaflets edges meet in the center of the valve, which we used as our focus point. With the model and the submersion box filled with the working fluid, we focused the lens and took an image of the calibration target, shown in Figure 3. Using the known distance of 2.6mm between each point on the calibration target, we calculated a conversion factor for our images of 0.00181mm^2^ per pixel. This results in approximately 580 pixels across the diameter of the valve, which allows us to accurately observe smallscale (*≈* 43*μm*) movements in the leaflets. Since the points on the target were in a Cartesian grid, we also used the target to reduce the distortion in the images caused by curved edges of the model with an in-house MATLAB code.

**Fig. 3.**
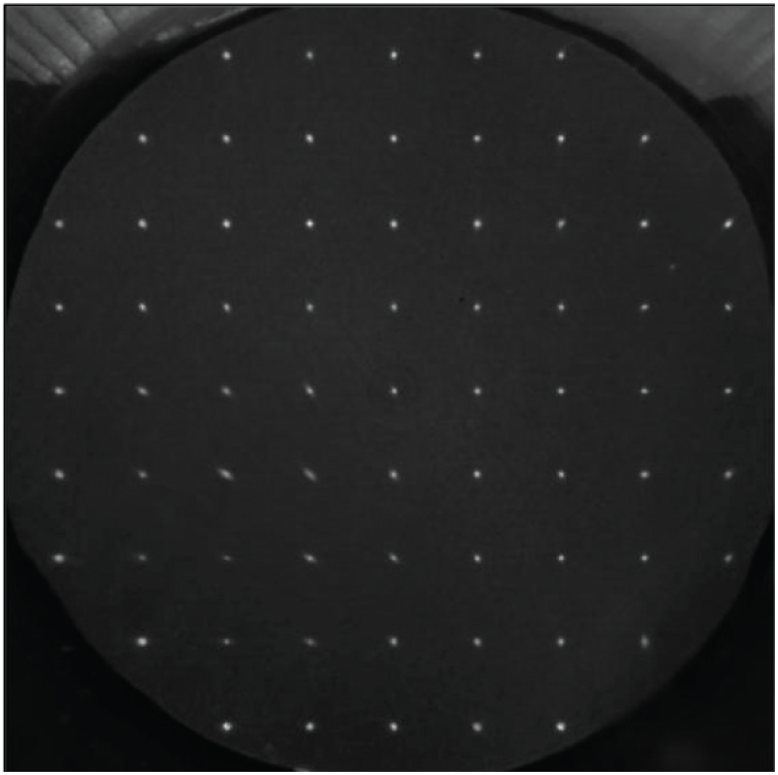
An image of the calibration target taken with the Phantom V2012 Ultrahigh-speed camera. The known distance between the grid point was used to calibrate the image pixel locations.

## 3 Results

First, we present results from the 4D-Flow MRI experiments. Then we examine valve leaflet motion with the high-speed imaging experiments and explore the relationship between leaflet motion and the velocity fields.

### 3.1 4D-Flow MRI

There were a number of similar flow features across the COs with the valve in the native orientation. In all three cases, the forward flow jet through the valve had an almost triangular shape with three tips, which was clearly seen in axial slice *x/D* = 0.5 at peak systole (Figure 4). Axial slice *x/D* = 0.5 is immediately downstream of the valve support structure. The jet generally followed the same angle and path out of the valve in all cases, as observed in the sagittal slice through the center of the vessel.

**Fig. 4.**
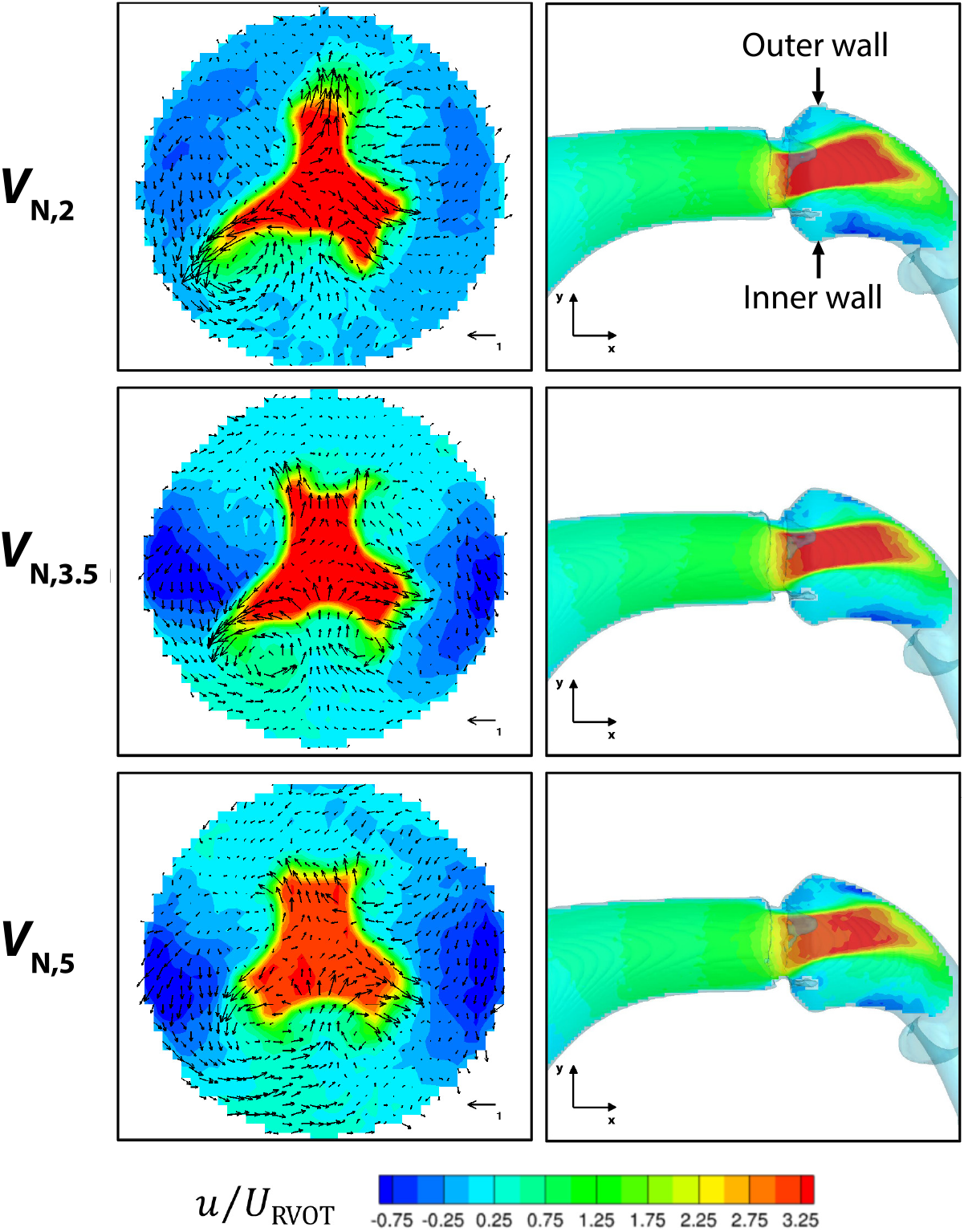
For COs of 2 L/min (top), 3.5 L/min (middle), and 5 L/min (bottom) with the valve in the native orientation, *x*-component of velocity contours normalized by the peak speed in the RVOT upstream of the valve. Contours shown at axial slice *x/D* = 0.5 (left column) and sagittal slice z = 0 (right column) at peak systole. In-plane velocity vectors are shown on the axial slice with a reference vector representing a normalized velocity of one.

In addition, the normalized streamwise velocities were similar for each CO, indicating that the relative strength of the jet compared to the inflow upstream of the valve was mostly independent of the actual CO.

However, despite these general similarities, there were key differences in the flow fields of each case. With a tri-leaflet valve, 120° symmetry would be expected in the flow through the valve, but this was not always the case. The forward jet shape was the most symmetric in *V*_N,5_, with each of the jet tips extending approximately the same distance away from the center of the vessel. *V*_N,3.5_ had more notable asymmetry as each of the jet tips had a unique shape, as observed in the axial slice *x/D* = 0.5. The jet asymmetry was the most pronounced in the 2L/min CO as demonstrated by the significantly elongates and narrowed jet tip, seen in Figure 4 in the bottom left of the *V*_N,2_ axial slice.

In *V*_N,2_, a strong vortex shed off the leaflet that bordered the narrowed asymmetric portion of the jet, as seen in the in-plane velocity vectors (Figure 4). There was recirculation present in a similar region in *V*_N,3.5_, but it was not as pronounced, and *V*_N,5_ only had limited recirculation surrounding the forward jet. The location and size of the reversed flow regions also differed across the three cases. In the sagittal slices, we can see a region of reversed flow along the interior curve in each case. In *V*_N,2_ this reversed flow formed fairly close to the valve annulus, while it was located further downstream in *V*_N,3.5_ and *V*_N,5_, with *V*_N,5_ having the smallest volume of reversed flow. However, there was a region of reversed flow above the forward jet in *V*_N,5_, and to a lesser extent in *V*_N,3.5_, that was not present in *V*_N,2_. These differences in the flow features depending on CO have clinical relevance, as increased recirculation and reversed flow have both been linked to valve complications such as calcification [23, 25].

The *x*-component of vorticity at peak systole illustrated additional flow features, as seen in contours on an axial slice that includes the valve leaflets and support structures at *x/D* = 0.25 (Figure 5). The *x* direction aligns with the streamwise flow and is normal to the valve annulus. For all the leaflets in all three CO cases, negative streamwise vorticity developed on one edge of the leaflet and positive streamwise vorticity developed on the other edge. These vortices aligned with the in-plane velocity vectors in each case. In *V*_N,2_, the stronger recirculation corresponded to a larger region of positive streamwise vorticity.*V*_N,5_ had weaker vorticity regions along the leaflets but had stronger vorticity on the outer edge of the valve supports, particularly along the outer curve of the vessel. This may indicate that the faster flow through the valve created a larger region of energized flow in the dilated anatomy. These variations in vorticity demonstrate how CO has a substantial impact on complex flow features in the RVOT model.

**Fig. 5.**
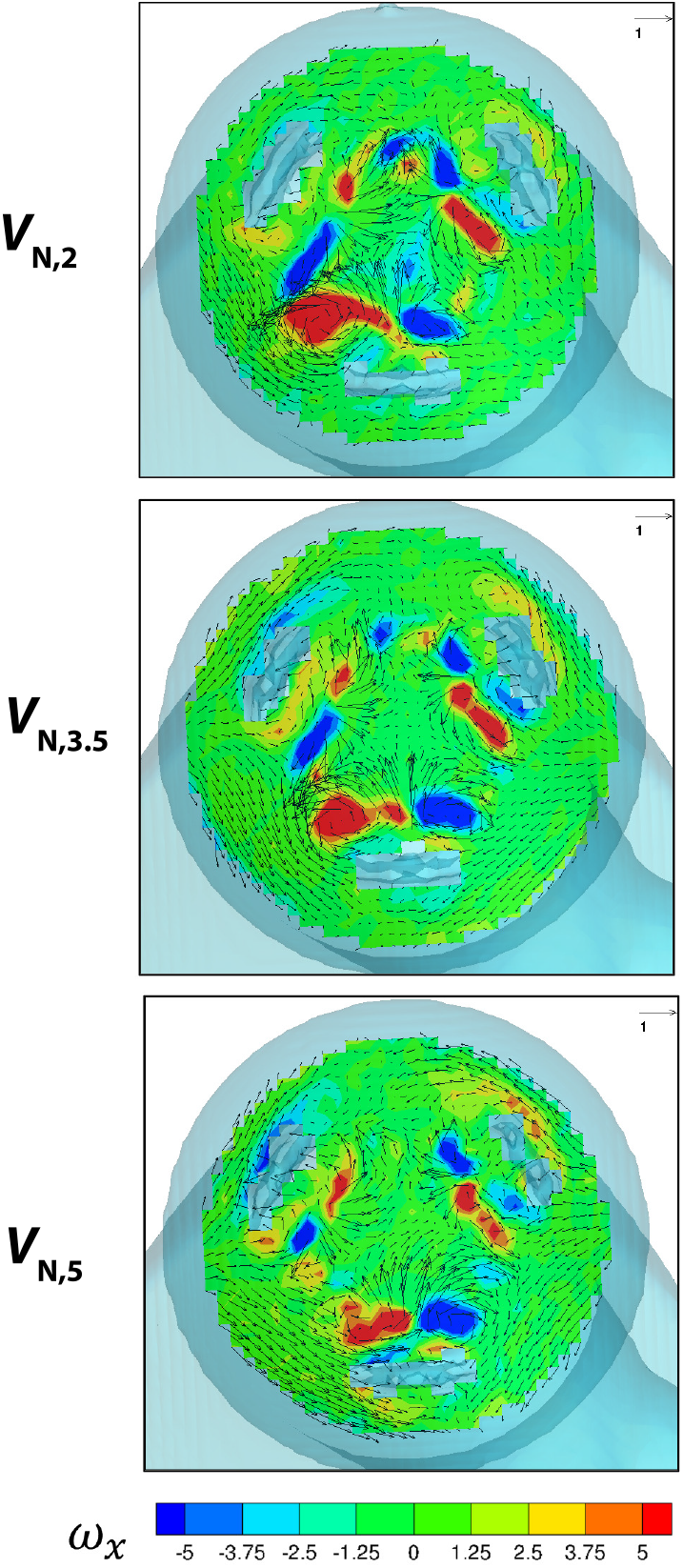
Contours of the x-component of vorticity (*ω*_*x*_) with in-plane velocity vectors at axial slice *x/D* = 0.25 for the 2 L/min (top), 3.5 L/min (middle), and 5 L/min (bottom) cases with native valve orientation at peak systole. The transparent light blue surface represents the wall of the model and the three structural supports of the valve, which cut through the axial contour slices.

Changing the valve orientation in addition to the CO had compound effects on the hemodynamics. In the rotated orientation, the valve was rotated 180° from the native orientation. The jet shape at systole, as observed in an axial slice, rotated 180° as well which can be seen when comparing the velocity contours for the native orientation in Figure 4 and the rotated orientation in Figure 6 for *V*_R,2_ and *V*_R,3.5_. In particular, *V*_R,2_ still had an asymmetric jet, with strong recirculation along the narrowed jet now along the outer wall of the vessel (upper right of the *V*_R,2_ axial slice in Figure 6). However, while the forward flow rotated accordingly with the valve orientation, the reversed flow regions at systole did not. In the native orientation, the large reversed flow regions formed mostly along the sides of the vessel. In the rotated orientation, the reversed flow regions were located along the inner and outer curves of the vessel. The sagittal slices illustrate the likely cause of this shift. In the rotated orientation, the jet through the valve annulus was directed towards the center of the vessel, away from the outer wall (Figure 6). This led to a large reversed flow region forming along that outer wall that was not present in the native orientation cases. Thus, changing the valve orientation drastically shifted the location and volume of reversed flow in both the 2 L/min and 3.5 L/min cases.

**Fig. 6.**
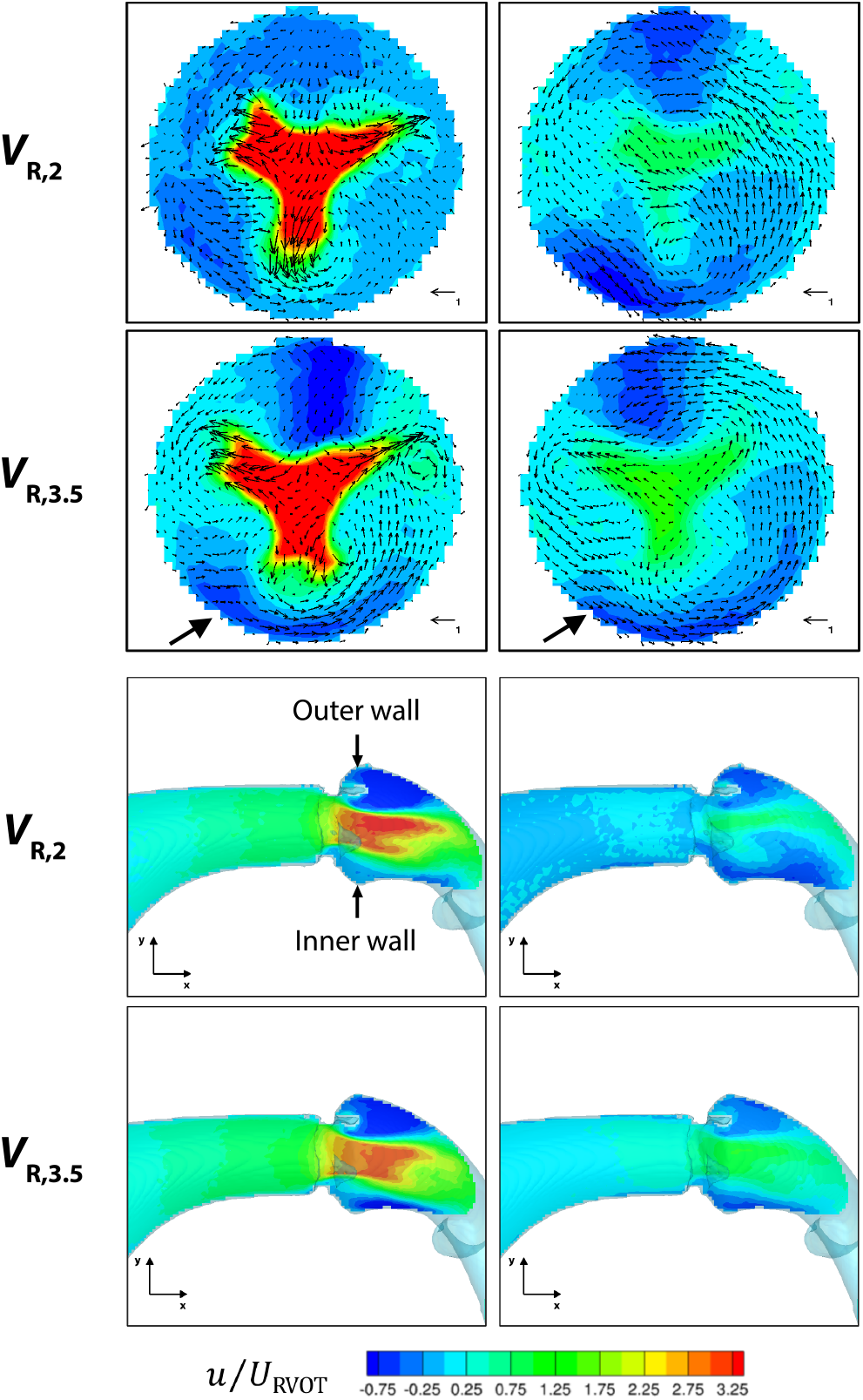
Normalized streamwise velocity contours for the 2 L/min (first and third rows) and 3.5 L/min cases (second and fourth rows) in the rotated valve orientation. The top block of four images are velocity contours shown at an axial slice *x/D* = 0.5 with in-plane velocity vectors. The bottom block of four images are velocity contours shown at a sagittal slice z = 0. Velocity contours are shown at peak systole and during diastole as flow is decelerating.

*V*_R,2_ presented a unique flow feature over the cardiac cycle that was not present in any other experiment in this study. In this case, a strong region of recirculation formed during systole along the inner curve of the vessel at one of the jet tips. This contrasted with *V*_R,3.5_, which did not have this recirculation feature. Instead, *V*_R,3.5_ had a strong counterclockwise swirling flow along the inner curve (indicated by the black arrow in the axial slice) that persisted into diastole. The recirculation in *V*_R,2_ appeared to break up that swirling effect that might have otherwise been present. As a result, the reversed flow along the outer curve, as observed in the sagittal slices, became detached from the wall in diastole for *V*_R,2_. This produced a more complex flow field in the center of the vessel around the leaflets, with reversed flow potentially impacting the point of valve closure. Increased recirculation and shifting reversed flow location may produce a hemodynamic environment more prone to leaflet calcification and fatigue that is only present in *V*_R,2_.

The radial velocities local to the valve directly impact the valve leaflets during opening and closure. Figures 7 and 8 demonstrate the radial velocities close to the valve leaflets at three different time points during the cycle: peak systole, the middle of diastole as the streamwise flow is decelerating, and towards the end of diastole when the streamwise flow has settled to rest. At systole, all CO and valve orientation cases experienced similar flow structures. Flow moved toward the center of the vessel between the open edges of the valve leaflets, as demonstrated by the blue velocity contours. Simultaneously, the opening surface of the leaflets pushed flow towards the outer wall of the vessel, resulting in the red radial velocity contours. The native and rotated orientations had similar flow structures, with the patterns rotated as they aligned with the location of the valve leaflets. However, after systole, the flows differed depending on both CO and valve orientation. During mid-diastole for the rotated valve orientation, both *V*_R,2_ and *V*_R,3.5_ had radial flow towards the vessel center mostly across the leaflet on the inner wall (the bottom of the axial slice) (Figure 8). This corresponded with the reversed flow regions seen in the axial and sagittal slices during diastole (Figure 6). Fluid was flowing back towards the valve annulus along both the inner and outer walls of the vessel. When it encountered the closed valve, it washed over the bottom leaflet towards the center of the vessel as radial flow. The radial flow on the top of the valve had significantly less momentum since it was blocked by the valve support structure. Towards the end of diastole, the radial flow became stagnant in both *V*_R,2_ and *V*_R,3.5_.

**Fig. 7.**
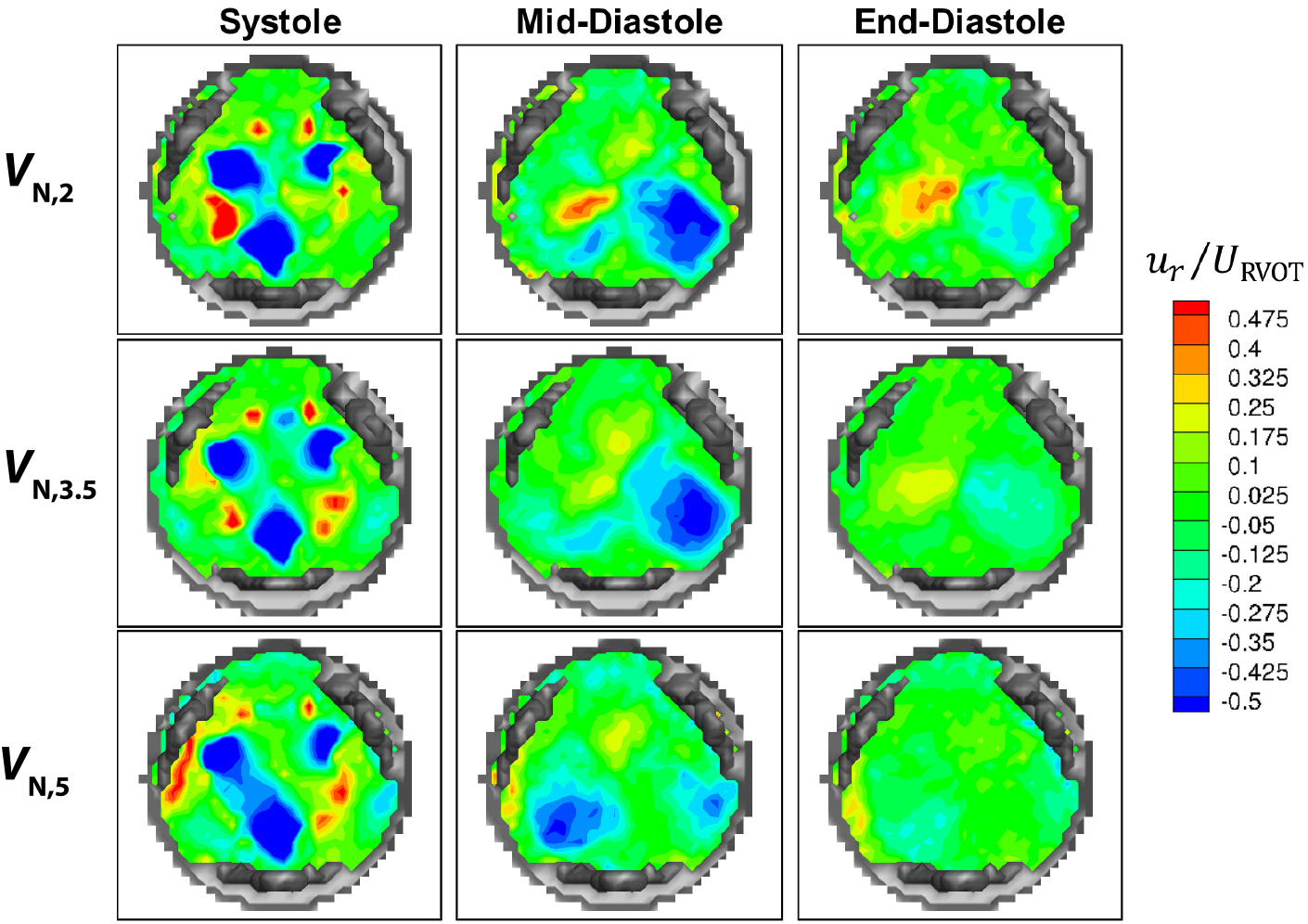
Radial velocity contours computed from the normalized velocity components for the native valve orientation for the 2 L/min case (top row), 3.5 L/min case (middle row), and 5 L/min case (bottom row). Contours shown at an axial slice *x/D* = 0.11, local to the valve annulus for three different time points: peak systole (left column), mid-diastole (middle column), and towards the end of diastole (right column). The structure of the valve is shown in gray.

**Fig. 8.**
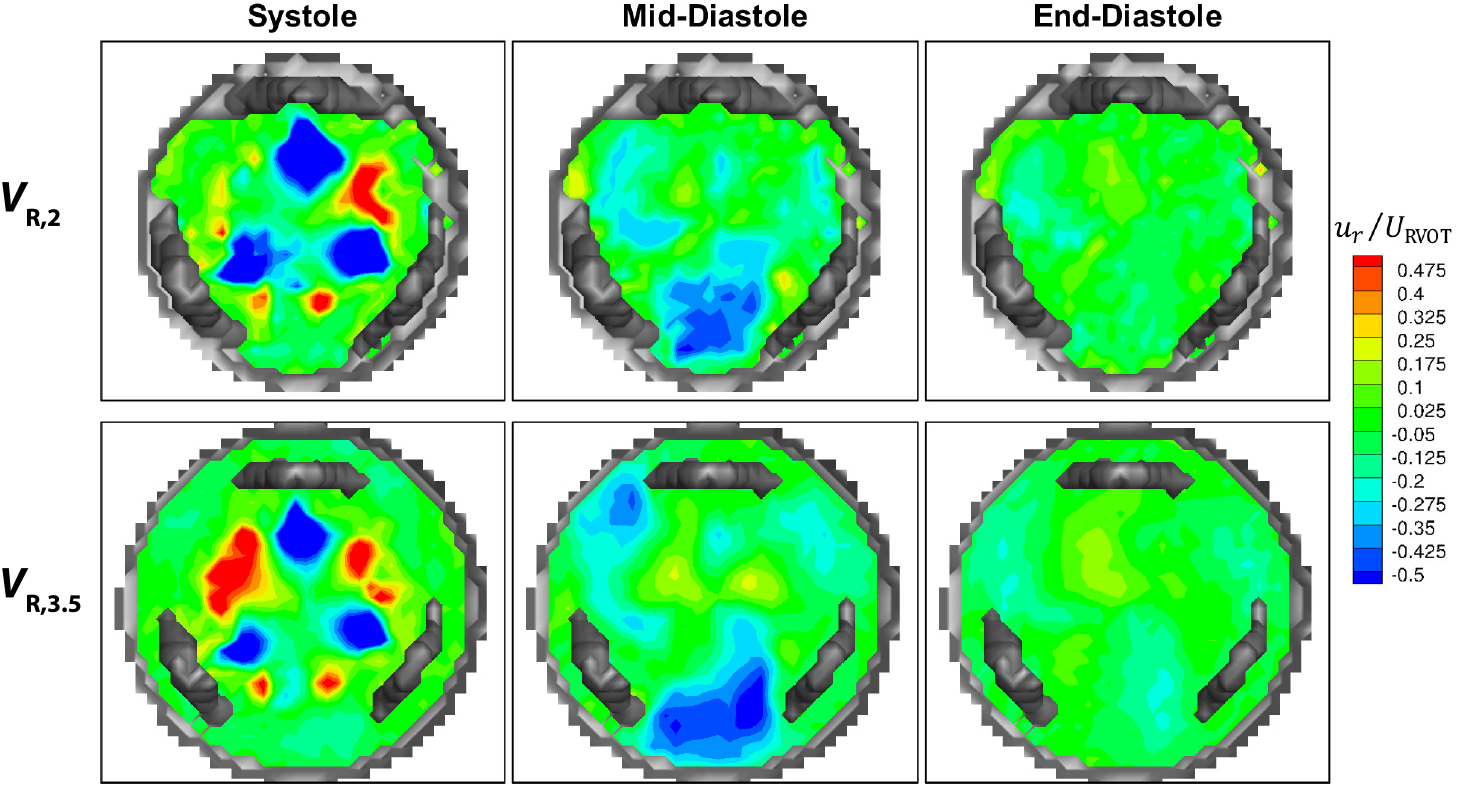
Radial velocity contours computed from the normalized velocity components for the rotated valve orientation for the 2 L/min case (top row), and 3.5 L/min case (bottom row). Contours shown at an axial slice *x/D* = 0.11, local to the valve annulus for three different time points: peak systole (left column), mid-diastole as the streamwise flow is decelerating (middle column), and towards the end of diastole when the streamwise flow has settled to rest (right column). The structure of the valve is shown in gray.

The radial flow patterns in diastole for the native orientation were substantially different than those for the rotated orientation. In *V*_N,2_ at mid-diastole, the radial flow moved across the closed valve from bottom right towards the center of the vessel (blue velocity contour) and then away from the center towards the bottom left region of the vessel (red velocity contour) (Figure 7). A similar flow pattern towards the center of the vessel was also seen in *V*_N,3.5_, though the radially outward flow was significantly weaker. In *V*_N,5_ at mid-diastole, flow moved towards the center of the vessel from both the bottom right and bottom left regions. Towards the end of diastole, when the streamwise flow had settled to rest, the radial flow in *V*_N,5_ was also stagnant. In *V*_N,3.5_, the flow was nearly stagnant, with some slight remnants of the flow patterns seen at mid-diastole. However, in *V*_N,2_, there was still a notable amount of radial flow washing over the valve leaflets. It is possible that larger volume of reversed flow led to this persistent radial flow throughout diastole, as more fluid flowed back towards the closed valve. This may have an adverse effect on valve performance, as continuous flow impacting the leaflets over the cardiac cycle may increase the rate of fatigue.

The cross-sectional slice *x/D* = 0.25 is approximately midway along the valve support structures and the valve leaflet edges extend to this point when they are fully open during systole. At this slice, the streamwise momentum *I*_1_ was approximately the same for the 3.5 L/min case with both orientations and the 5 L/min case (Figure 9). The streamwise momentum for the 5 L/min case peaked at an earlier phase because that was when peak systole occurred for the 5 L/min case flow rate waveform. Otherwise, the *I*_1_ values for these cases were very similar, with *I*_1_ exceeding 1 during systole indicating high momentum flow and decreasing to 0 when the streamwise flow stagnates during diastole. Both *V*_N,2_ and *V*_R,2_ followed a similar pattern over the cardiac cycle, but peaked at a higher value of *I*_1_ during systole. One possible explanation for the higher streamwise momentum in the 2 L/min cases is that the valve leaflets did not open as fully due to the slower CO through the 25mm valve. The flow through the narrowed valve opening would be faster and result in higher relative streamwise momentum in this case. We observed that the valve opening area was less for the 2 L/min cases than the others with the high-speed imaging, as discussed later, indicating that this was likely the cause of the higher streamwise momentum in these cases.

**Fig. 9.**
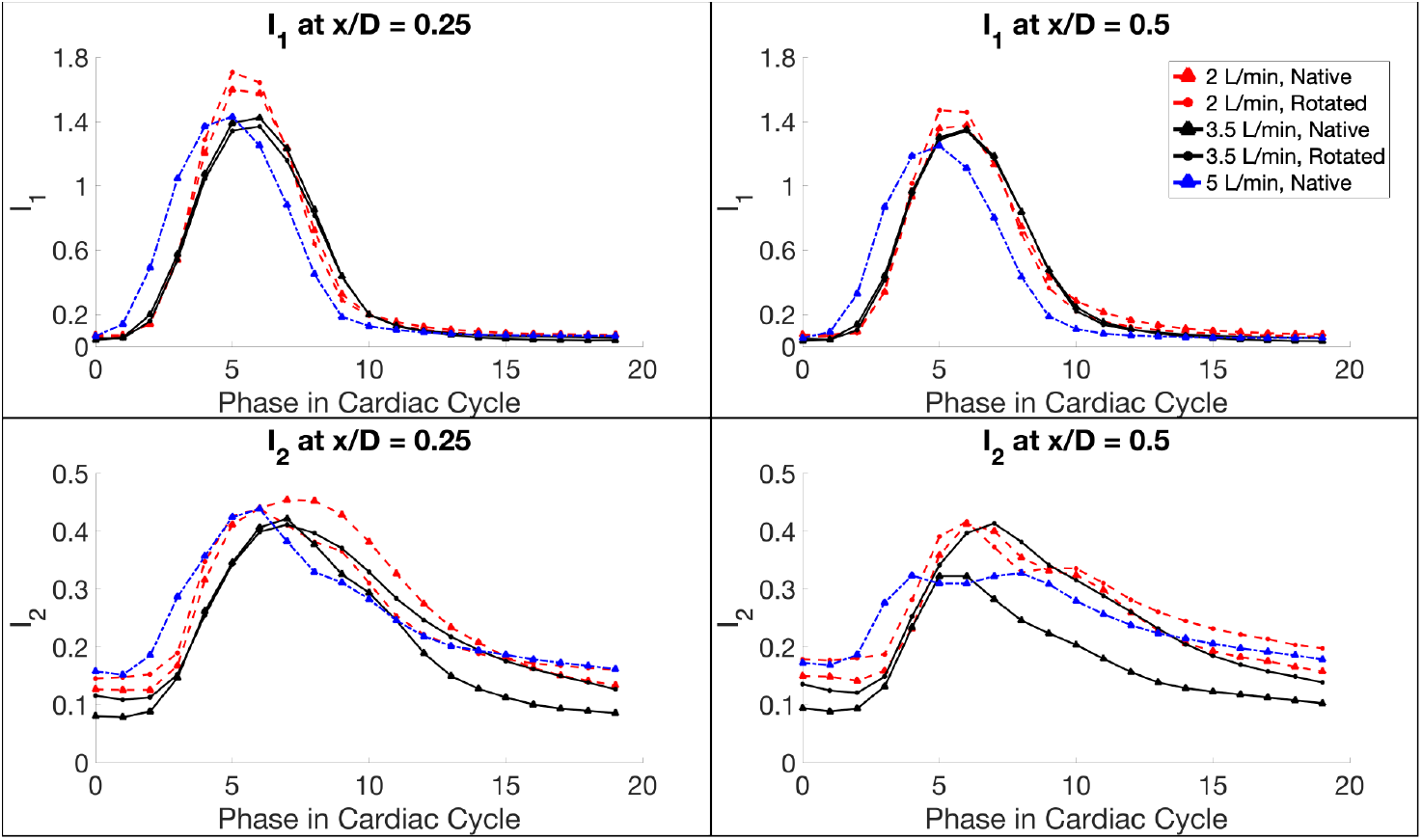
Quantitative integral metrics *I*_1_ (top row) and *I*_2_ (bottom row) at two axial slices, *x/D* = 0.25 (left column) and *x/D* = 0.5 (right column) for three COs. The legend in the top right corner applies to all plots.

The cross-sectional slice *x/D* = 0.5 is immediately downstream of the entire valve support structure. At this slice, *I*_1_ had similar patterns and relationships across cases as *I*_1_ at *x/D* = 0.25 (Figure 9). For all cases, *I*_1_ was decreased at *x/D* = 0.5 as the jet spread and slowed moving downstream. *I*_1_ for the 2 L/min cases was still slightly higher than the other cases implying that the valve’s orifice area is significantly lower for this case. For all cases at this slice, the streamwise momentum peaked above 1 during systole and decreased to 0 during diastole. For both *x/D* = 0.25 and *x/D* = 0.5, there were only slight differences between the native and the rotated orientation for the 2 L/min and 3.5 L/min cases, indicating that valve orientation had little impact on the streamwise momentum.

For the secondary flow strength *I*_2_ at *x/D* = 0.25, all of the cases had similar peak values, with the 3.5 L/min cases on the lower side at 0.4 and *V*_N,2_ at the highest value of 0.45 (Figure 9). In *V*_N,2_, *I*_2_ was sustained close to its peak value for longer in the cardiac cycle than the other cases. This higher *I*_2_ value with longer duration corresponded to larger region of *x* vorticity that occurred in this case (Figure 5). *V*_N,2_ had higher *I*_2_ after systole than *V*_R,2_, while the opposite was true for the 3.5 L/min cases. This highlights the compound influence of CO and valve orientation; the effect of changing the valve orientation depends on the flow rate. At *x/D* = 0.5, both 2 L/min cases and the *V*_R,3.5_ had nearly the same *I*_2_. For *V*_N,5_ at *x/D* = 0.5, there is no clear peak, but rather a consistent secondary flow strength over the entirety of systole. *V*_R,3.5_ had a strong counterclockwise swirl at this axial location, differentiating it from *V*_N,3.5_ (Figure 6). The higher value of secondary flow strength quantified this difference. Both of the 2 L/min cases had strong recirculation regions along the forward flow jet which contributed to the higher *I*_2_ values. It is worth noting that the differences in *I*_2_ at the beginning and end of the cardiac cycle, when the flow is essentially stagnant, are likely due to differences in the noise floors in the secondary flows across cases as opposed to physical differences in the flows.

### 3.2 High-Speed Imaging

We analyzed valve leaflet behavior with high-speed imaging experiments for all three COs and two valve orientations for each case. From the 2D images of the valve, we calculated the orifice area over the cardiac cycle. For each case, we captured images for 48 cardiac cycles, calculated the projected 2D orifice area in each image by counting the pixels inside the leaflet edges using custom in-house MATLAB code, and averaged the area over all cardiac cycles (Figure 10). The orientation of the valve did not significantly impact the maximum orifice area during peak systole, though it did have slight effects on the timing of valve opening and closure. The 5 L/min cases had the highest orifice area, indicating that these cases made the most efficient use of the valve. The 3.5 L/min cases had a 4.5% lower orifice area and the 2 L/min cases had a 8.9% lower orifice area than the 5 L/min cases. With the same 25mm valve, the lower COs were not sufficient to fully open the leaflets.

**Fig. 10.**
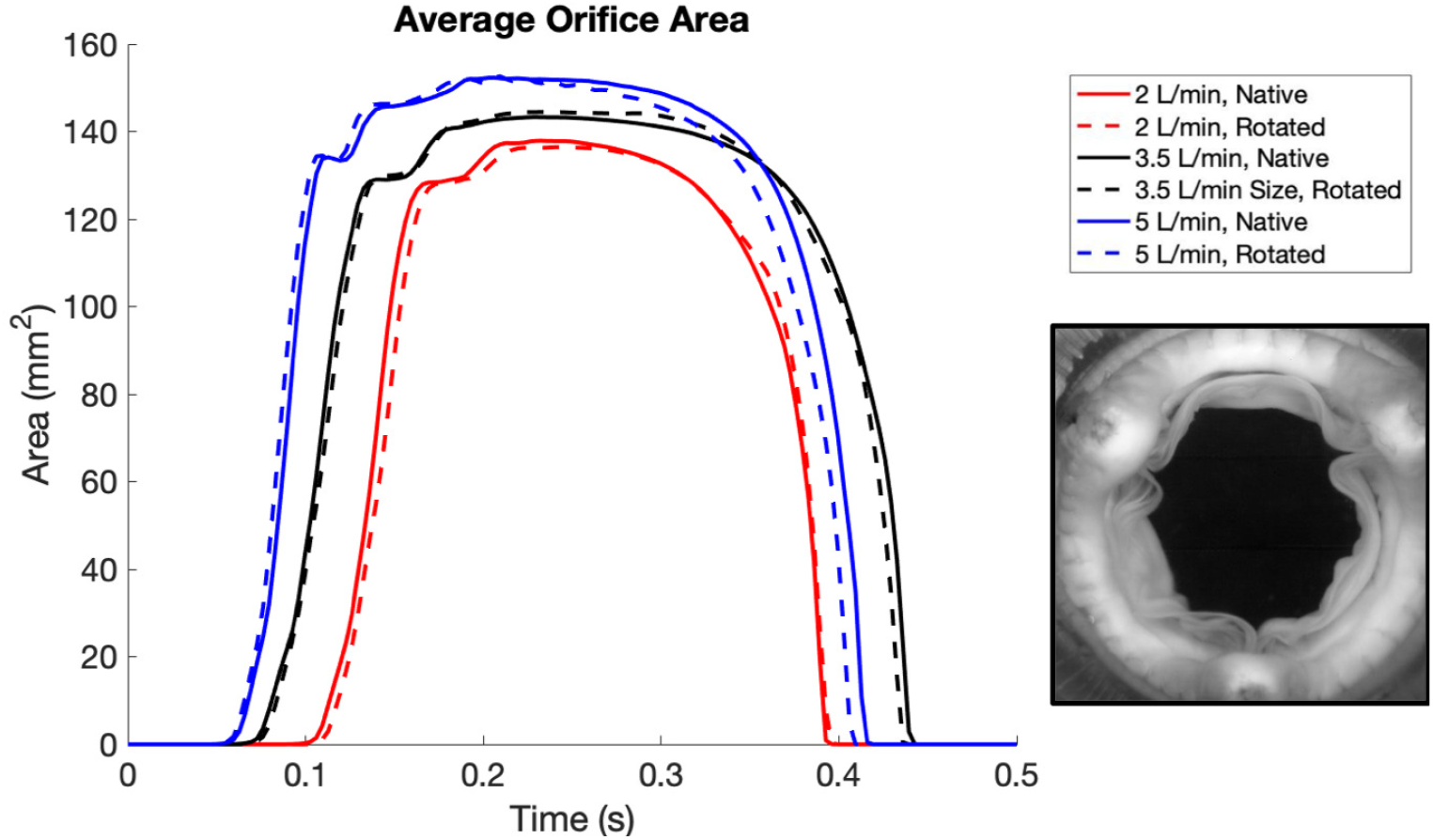
Average orifice area for three COs and two valve orientations over the systolic portion of the cardiac cycle. Orifice area was calculated as the projected area between the open valve leaflets in the 2D valve image (example shown in bottom right inset).The image shows the valve leaflets with the valve in the native orientation in the 3.5 L/min case.

While the leaflet motion varied slightly from cycle to cycle, there were some key features of the leaflet behavior that differentiated the cases. For example, two critical differences between *V*_N,2_ and *V*_N,3.5_ are highlighted in the white boxes in Figure 11. For these regions, *V*_N,5_ was qualitatively similar to *V*_N,3.5_. The top white box focuses on an area where the leaflet free edges meet at the commissure. In *V*_N,3.5_, the leaflet edges fully separated, pushed open by the incoming flow. However, in *V*_N,2_, the leaflet edges remained in contact through the cardiac cycle. This contact can adversely affect the valve performance, as leaflet edges close together at the commissure may calcify more easily, leading to valve dysfunction. In the bottom white boxes, we observed a difference in the shape of one of the leaflets. While this section of leaflet did not fully open to the outside of the valve in *V*_N,3.5_, it came closer than it did in *V*_N,2_. With a CO of 2 L/min, this section of leaflet remained in an open bowl-like shape, which obstructed the flow coming through the annulus and provided an area where flow could recirculate and stagnate, even during systole.

**Fig. 11.**
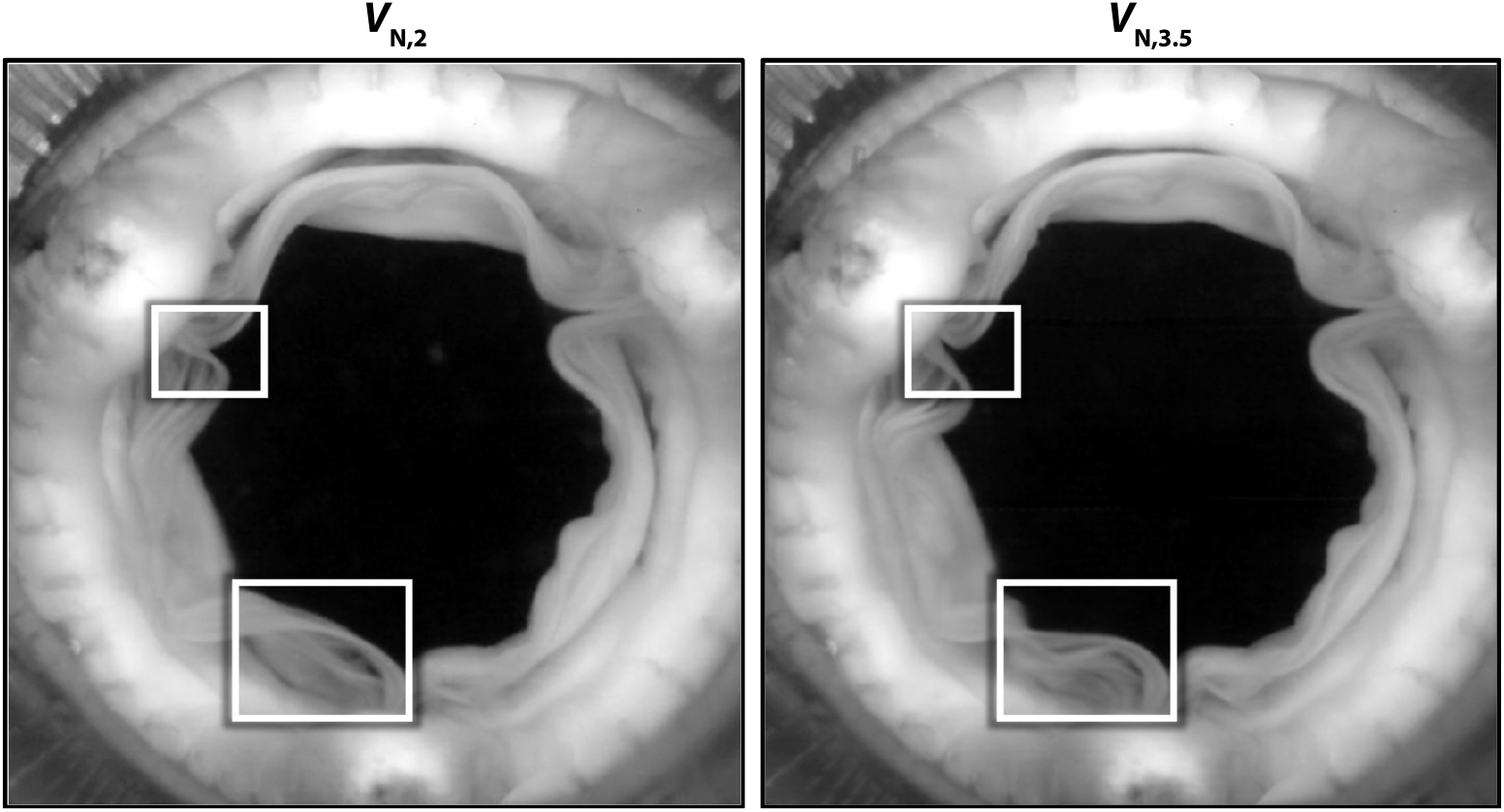
Valve leaflet images from the high-speed imaging experiments for the 2 L/min case (left) and the 3.5 L/min case (right). The white boxes highlight differences in leaflet behavior between the two cases.

These observations of the valve leaflet behavior are useful on their own, as they demonstrate which cases result in valve motion that might be more prone to dysfunction. However, we also can couple the high-speed imaging with our 4D-Flow MRI results to fully examine the relationship between valve behavior and flow patterns. Key examples of this relationship are shown for *V*_N,2_ and *V*_R,2_ at slice *x/D* = 0.5 at peak systole in Figure 12. For the native case, the leaflet that we observed obstructing the flow in Figure 11 directly corresponded to the region of strong recirculation in the 4D-Flow MRI data. The flow from the 2 L/min CO was not fast enough to fully open the leaflet, which led to adverse hemodynamics in the velocity field. This behavior also occurred in *V*_R,2_. The two white boxes in the bottom row of Figure 12 show two areas where valve leaflets did not open fully. This directly corresponded to the narrowed jet region in the velocity data, as well as another region of recirculation. These hemodynamic features can contribute to increased risk of calcification and uneven leaflet fatigue due to asymmetry.

**Fig. 12.**
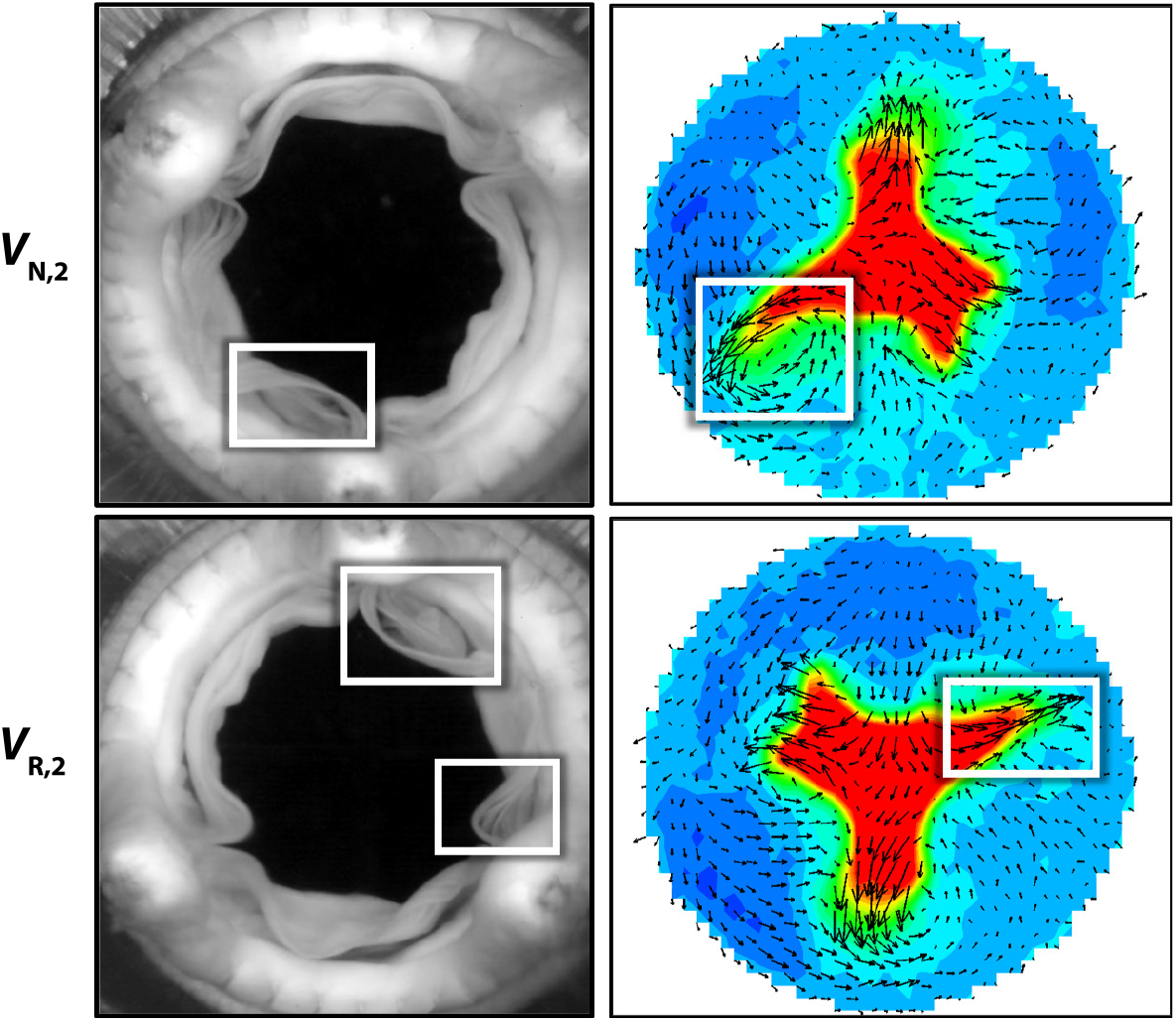
Valve leaflet images compared to velocity contours for the 2 L/min case with the native valve orientation (top row) and rotated valve orientation (bottom row). The white boxes highlight regions where the leaflet behavior observed in the high-speed imaging experiments align with strong recirculation in the velocity fields obtained by 4D-Flow MRI at slice *x/D* = 0.5 at peak systole.

## 4 Conclusions

The analysis of full 4D-Flow fields and instantaneous valve leaflet images demonstrated how both CO (which is a proxy for valve sizing in our experiment) and valve orientation can impact hemodynamics in the RVOT and leaflet performance. The general shape of the jet through the valve and the relative velocity of the jet compared to the upstream inflow were similar across COs of 2 L/min, 3.5 L/min, and 5 L/min. However, there were key differences in the asymmetry of the jet, the amount of recirculation, and the location and size of reversed flow regions (Figure 4). The 5 L/min case had the most symmetric jet and the lowest volume of reversed flow. Conversely, the 2 L/min case had an asymmetric jet with a strong region of recirculation along the narrowed branch of the jet. This recirculation corresponded to a larger region of positive *x* vorticity in the 2 L/min case, though all three cases had similar vortical structures as two vortices of opposite sign shed off of each valve leaflet (Figure 5).

In *V*_R,2_ and *V*_R,3.5_, the forward flow jet rotated 180°, following the valve rotation. However,the reversed flow regions drastically differed from the native orientation cases. In particular, *V*_R,2_ had increased recirculation along the inner curve of the vessel, which contributed to the reversed flow detaching from the wall and impacting the valve leaflets during diastole. These unique flow features emphasized the compound effect of CO and valve orientation. Both of these factors can have a crucial impact on hemodynamics and trends observed from changing valve orientation in one case were not necessarily replicated with a different CO. These compound effects were also quantified in the integral metric *I*_2_ (Figure 9). At axial slice *x/D* = 0.5, both *V*_N,2_ and *V*_R,2_ had similar secondary flow strengths, while *V*_R,3.5_ had substantially higher secondary flow strength than *V*_N,3.5_. Thus, the effect of valve orientation on secondary flow strength depended on the CO.

High-speed imaging experiments provided insights on the valve leaflet behavior with different COs. We used the images to measure the valve orifice area and noted that the 2 L/min cases had 8.9% lower area than the 5 L/min cases. Orifice area is used extensively in clinical settings, particularly with aortic valves, to assess performance of prosthetic valves; lower orifice area indicates a less efficient valve [43–45]. In addition to the orifice area calculations, the combination of our 4D-Flow MRI and high-speed imaging methods provided a more complete picture of the valve environment. In several instances with the 2 L/min cases, we observed leaflet sections that did not open fully during systole directly corresponded to areas of strong recirculation in the velocity data. These connections, in addition to providing insight on valve performance, indicate that it is possible to infer valve leaflet behavior from flow fields. This relationship would be useful for *in vivo* settings where it is possible to obtain velocity data, but not possible to directly observe leaflet motion.

We found that the 2 L/min cases had numerous adverse features in the flow fields and valve behavior. The 2 L/min cases had strong recirculation, which is associated with calcification, large reversed flow volumes, which may lead to hemolysis, persistent radial flow during diastole, which may contribute to leaflet fatigue, lower valve orifice area, and locations where the valve leaflet edges did not separate, which are more prone to fusion and calcification [23–25]. As discussed earlier, a CO of 2 L/min through a 25mm valve corresponds to the clinical decision of valve oversizing. While the effects of this practice are still an open question, our results strongly suggest that valve oversizing produces a hemodynamic environment that would predispose the valve for failure. Alternatively, we found that the 5 L/min case, which corresponds to valve undersizing, had fewer adverse flow features and the highest orifice area in our cases. While there are a number of clinical reasons to avoid undersizing the valve, including needing to account for patient growth, our results indicate that the hemodynamics of this configuration are comparable or even favorable to the standard sizing.

While this study provided many key insights about the effect of valve sizing and orientation on RVOT hemodynamics, clinical studies are necessary to determine how flow features are correlated with adverse outcomes from PVR surgery. Clinical studies could ascertain the association between flow features, such as recirculation, and long-term outcomes, thus establishing which hemodynamic environments are the most favorable for valve function. In addition, we used the same valve for all of our experiments for consistency. However, it is possible that some of the features we observed, particularly in the high-speed imaging experiments, might be a property of this specific valve as opposed to all bioprosthetic valves.

Overall, we demonstrated the compound effect of valve sizing and orientation on RVOT hemodynamics and valve performance. The combination of 4D-Flow MRI and high-speed imaging allowed us to obtain a full view of the valve environment. In particular, we determined that the 2 L/min case had multiple features in the flow fields and valve leaflet behavior that could adversely affect long-term valve performance.

## Acknowledgements

The authors thank their funding sources: Stanford’s Maternal and Child Health Research Institute, the American Heart Association, the Stanford Bio-X Bowes Fellowship, and the National Science Foundation Graduate Research Fellowship.

## Declarations

N.K. Schiavone, P.J. Nair, C.J. Elkins, D.B. Ennis, J.K. Eaton, and A.L. Marsden declare they have no conflicts of interest. D.B. McElhinney is a proctor and consultant for Medtronic, which is not directly relevant to this study.

